# Prolific non-research authors in high impact scientific journals: meta-research study

**DOI:** 10.1101/2022.11.19.517227

**Authors:** John P.A. Ioannidis

## Abstract

Journalistic papers published in high impact journals can be very influential, especially in hot fields. This meta-research analysis aimed to evaluate the publication profiles, impact, and disclosures of conflicts of interest of non-research authors who had published >200 Scopus-indexed papers in Nature, Science, PNAS, Cell, BMJ, Lancet, JAMA or New England Journal of Medicine. 154 prolific authors were identified, 148 of whom had published 67,825 papers in their main affiliated journal in a non-researcher capacity. Of 25 massively prolific authors with over 700 publications in one of these journals, only 3 had a PhD degree in any subject matter. Only 2 of the 25 disclosed potential conflicts with some specificity. The practice of assigning so much power to non-researchers in shaping scientific discourse should be further debated and disclosures of potential conflicts of interest should be emphasized.

## INTRODUCTION

Publications in high impact peer-reviewed scientific journals are influential and fervently coveted (Hammarfelt, 2017). For many scientists, even a single publication in such venues can represent a unique career achievement. Researchers struggle to publish with low single-digit acceptance percentage rates (Herbert, 2020) and painstaking peer review. However, some authors readily publish several hundreds or even more than a thousand publications in these same journals. These prolific authors are editorial staff and science writers who write routinely for a journal on diverse matters, ranging from editorial opinions to news items, perspectives, features, and/or surveys. Their writings can be published expediently (sometimes even within hours of submission) and without formal peer review. The exact acceptance rates are unknown, but are probably very high. Appearing in venues of very high visibility, these writings may yield major influence on science, scientific debate, science policy, and the public perception of science. Moreover, given their non-technical nature they can reach wider audiences than technical papers written by researchers and the importance of sound, balanced, and accurate non-technical science communication cannot be overstated (Trese & Weigold, 2002; Bublea et al., 2009)., Concurrently, given the central role that these elite science writers can play in both science and policy, it would be essential to have transparent information on their potential conflicts of interest.

To our knowledge, there has been no prior systematic bibliometric evaluation of prolific non-research authors publishing in high impact journals. Given the potentially vast influence they can exert, it is essential to study this phenomenon. The current evaluation provides a systematic mapping and analysis of prolific authors who have published more than 200 papers in at least one high impact scientific journal of general science or medicine; and a more in-depth analysis of those who are massively prolific and have published over 700 papers in at least one such high impact scientific journal. It aims to characterize the productivity record of these science writers and its impact, as well as their credential profile and conflict of interest disclosures.

## METHODS

### Eligible journals and prolific authors

The current analysis focused on 4 general science journals (Nature, Science, Proceedings of the National Academy of Science USA [PNAS], and Cell) and 4 general medical journals (New England Journal of Medicine [NEJM], Lancet, Journal of the American Medical Association [JAMA], and British Medical Journal [BMJ] that are considered to be highly prestigious and highly desirable and extremely competitive publication venues for scientists. Moreover, the analysis focused on authors who have published in their career more than 200 Scopus-indexed publications in one of these journals, regardless of the total number of their publications. Authors who published more than 200 publications in these 8 journals combined but did not have >200 publications in at least one of them were not eligible. The threshold of 200 was pre-specified in an arbitrary way, aiming to ensure that only few, if any, of the retrieved authors would represent academic researchers without editorial/journalistic roles who manage to publish an extreme number of papers in a single journal. It also aimed to capture the most productive, and thus potentially most influential, among the journalistic authors penning articles in/for these journals. Authors were eligible regardless of whether they belonged officially to the journal staff or were free lancers.

### Data sources and search strategy

The Scopus database (Baas, et al., 2020) was used to search for eligible authors and their publication corpora. Searches were last updated on August 26, 2022. For each of the eligible journals, all items published and indexed in Scopus were retrieved using the SRCTITLE command (source title searches). For BMJ and JAMA different writings of their name were included (BMJ, The BMJ, BMJ (online), British Medical Journal, British Medical Journal clinical edition; Journal of the American Medical Association, and JAMA). The Scopus search engine shows the most prolific authors in decreasing order of number of publications. All authors with an author ID file including more than 200 published items were considered eligible.

### Data extraction

For each journal, the following information was extracted: total number of published items, number of authors with >200 published items in this journal, total number of published items authored by these prolific authors, number of published items authored by these prolific authors that had received more than 100 citations, and the respective number that had received more than 20 citations.

The eligible prolific authors were also assessed on whether they might be publishing predominantly as researchers rather than with primarily non-researcher (editorial, staff writer, invited/freelance science writer, correspondent, news writer) roles in the journal. In the main analyses, full-time academics publishing predominantly as researchers were excluded from the analysis of total published items by prolific authors and their citation impact.

The published items by the prolific non-researcher authors were also evaluated for their categorization/classification by Scopus. Scopus assigns the published items in the following categories: editorial, note, article, short survey, letter, conference proceeding, review, and erratum.

The recently published (2020-2022) publications that had already received over 100 citations were examined in-depth for the types of topics that they covered.

### In-depth evaluation of the most massively prolific authors

For the most massively prolific authors, i.e. those who had published more than 700 publications in a single high impact journal, focused searches were performed trying to identify all their publications, regardless of what journal these publications had appeared in. These searches used the last and first name of the author and in some occasions they identified some additional Scopus author ID files that belonged to the same author. Precision and recall in Scopus are very high (98% and 93.5%) (Schotten et al., 2017) which means that almost all publications by a given author are in a single author ID file in Scopus and almost all publications clustered in a given author ID are by the same author. However, for some prolific authors it is possible that a minority of their publications are split in separate smaller author ID files. Here, these split files were merged to obtain a complete picture of the productivity of each eligible author. Then, it was possible to obtain the number of citations received by all their publications combined, the number of citations received by their publications in their main affiliated journal, the number of citations by their most highly-cited publication published in a journal other than their main affiliated journal, and the number of citations received by their most highly-cited publication that was not a journalistic/editorial/news piece, but a research paper (either primary research or secondary research, e.g. a formal review or guideline).

Finally, Google searches with the name of each author tried to identify whether there is any readily retrievable information on their education credentials – specifically, PhD or equivalent degrees on any scientific field, MD, and Master’s degree in journalism topic. Moreover, the website of the journals where they published prolifically was screened to identify if any information was listed on conflicts of interest disclosures.

## RESULTS

### Eligible prolific authors and their publication corpora

Table 1 shows the eligible prolific authors, their total publication corpora and the citation impact thereof for each of the 8 examined journals. As shown, Cell did not have any so prolific authors meeting the definition of eligibility. The most prolific author in Cell, a Nobel laureate, had published 82 items therein during his career. Conversely, the other 7 journals had anywhere from 3 to 43 eligible prolific authors each, for a total amounting to 154. Of the 154, one was anonymous (Lancet Editorial). All three prolific authors of PNAS were publishing in their capacity as researchers and the same applied to 1 prolific author in JAMA and 2 prolific authors in BMJ (although not all of their published papers represented research work, several were still of editorial nature). Excluding these 6, the remaining 148 author profiles reflected writers who acted on non-researcher (editorial, staff writer, invited/freelance science journalist, correspondent, news writer) roles in their affiliated journal. The 148 profiles pertained to 146 different authors (two individuals had been prolific in two of these journals each). Of note, 7 of the 8 prolific authors of the England Journal of Medicine were involved in editing the highly popular weekly Case Record series; they are typically not listed as authors in PubMed (other scientists were listed as traditional authors for the same articles). Another 7 prolific authors were editors-in-chief or senior editors. All the remaining were journalistic authors, who may have had a formal role in the journal (e.g. listed in the news section in-house staff writers) or be entirely freelance.

**Table 1.**
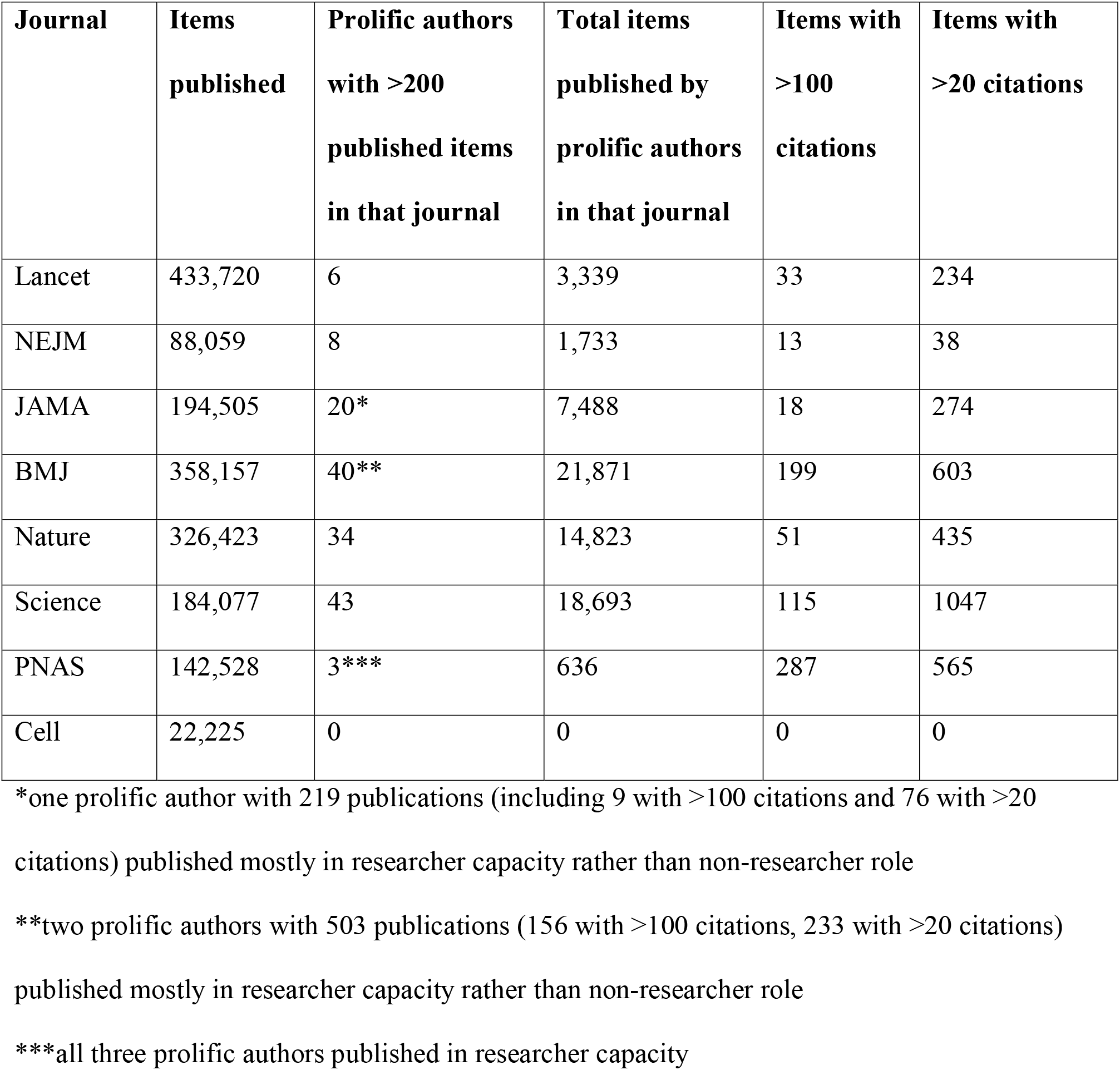
Publications by prolific authors and their impact in the eligible high impact journals.

| **Journal** | **Items published** | **Prolific authors with >200 published items in that journal** | **Total items published by prolific authors in that journal** | **Items with >100 citations** | **Items with >20 citations** |
| --- | --- | --- | --- | --- | --- |
| Lancet | 433,720 | 6 | 3,339 | 33 | 234 |
| NEJM | 88,059 | 8 | 1,733 | 13 | 38 |
| JAMA | 194,505 | 20* | 7,488 | 18 | 274 |
| BMJ | 358,157 | 40** | 21,871 | 199 | 603 |
| Nature | 326,423 | 34 | 14,823 | 51 | 435 |
| Science | 184,077 | 43 | 18,693 | 115 | 1047 |
| PNAS | 142,528 | 3*** | 636 | 287 | 565 |
| Cell | 22,225 | 0 | 0 | 0 | 0 |

\*one prolific author with 219 publications (including 9 with >100 citations and 76 with >20
citations) published mostly in researcher capacity rather than non-researcher role
\*\*two prolific authors with 503 publications (156 with >100 citations, 233 with >20 citations)
published mostly in researcher capacity rather than non-researcher role
\*\*\*all three prolific authors published in researcher capacity

The prolific authors had a total of 68,643 items published in their keys affiliated journals alone (67,285 based on those with non-researcher roles), and this may be a modest underestimate given that only their main author ID Scopus files were considered in this calculation (see also below). A total of 716 published items (1.0%) had received more than 100 citations in Scopus. The number dropped to 264 (0.4%) when the 6 researcher authors were excluded. A total of 3,196 published items (4.7%) had received more than 20 citations in Scopus. The number dropped to 2,322 (3.5%) when the 6 researcher authors were excluded.

### Types of published items per Scopus classification (Table 2)

**Table 2.**
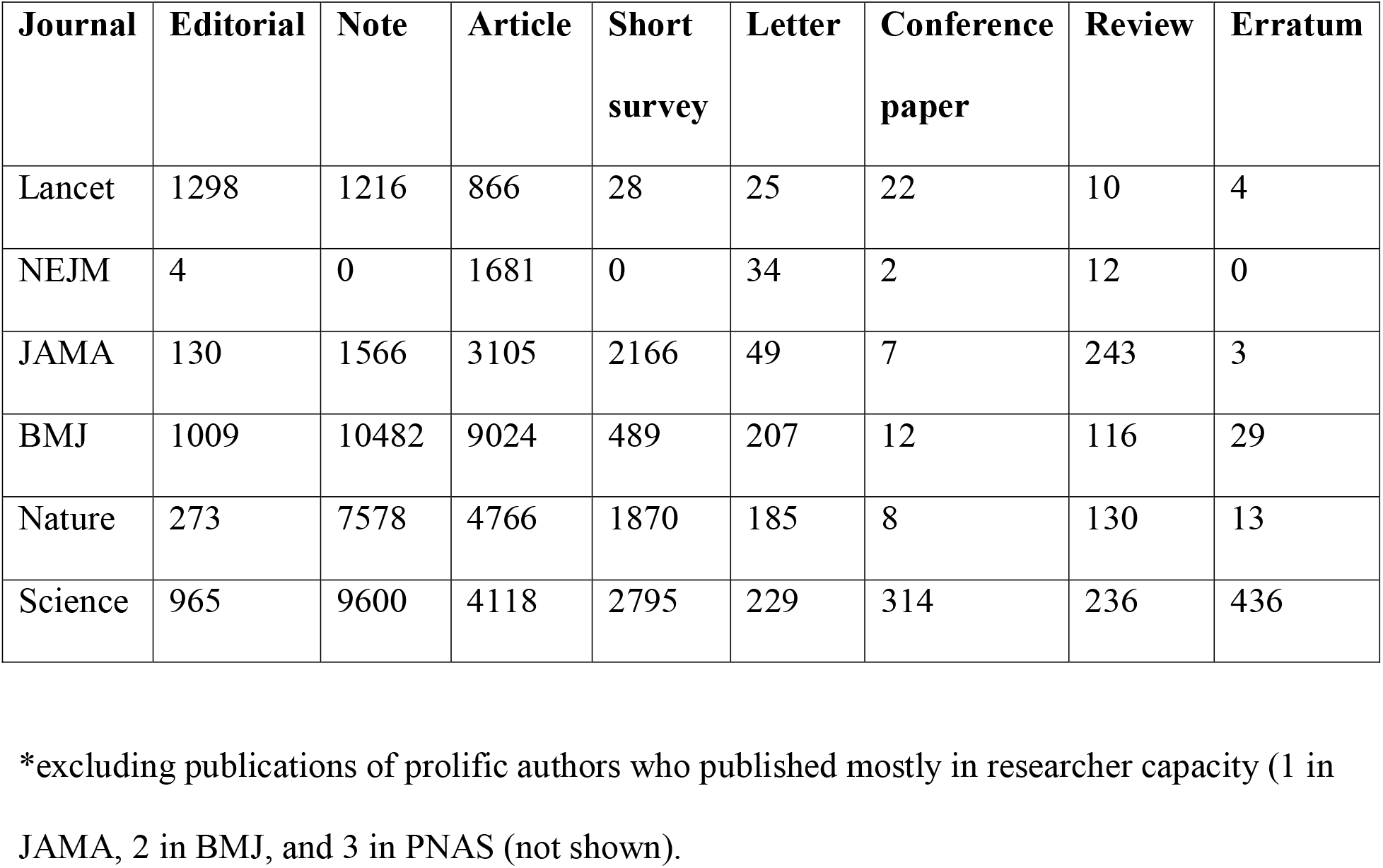
Categorization by Scopus of the publications of the prolific authors*.

Among the 67,285 items published by prolific authors with non-researcher roles, Scopus characterized as “editorials” only a small minority (3,679, 5%). The most common characterizations were as “articles” (23,560, 35%), or “notes” (30,442, 45%), and many other publications were characterized as “short surveys” (7,348, 11%). “Letters” and “reviews” accounted for approximately 1% each and there were even smaller numbers of items characterized as “conference papers” and “errata”. There were differences across journals on what were the most common characterizations. “Notes” were the most common characterization in 4 journals, but were absent in NEJM and less common in JAMA. In the latter journals, the most common characterization of the published items was as “articles”.

### Recent highly-cited papers by prolific authors

41 papers published in 2020-2022 by the prolific authors in their affiliated journals had already received >100 citations (Table 3). 40 of them were on hot (and often controversial), rapidly emerging areas of COVID-19 and one was on artificial intelligence.

**Table 3.**
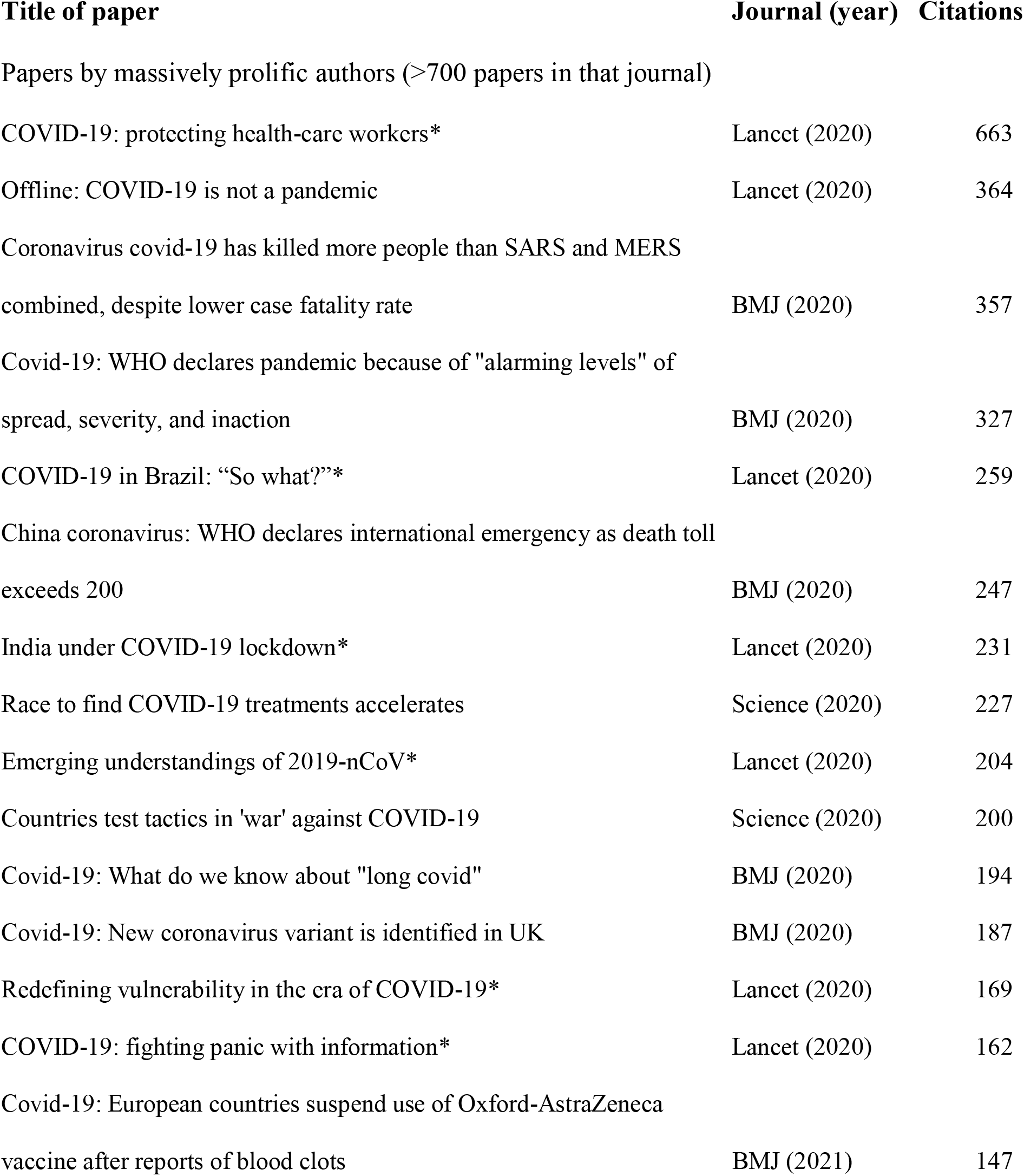
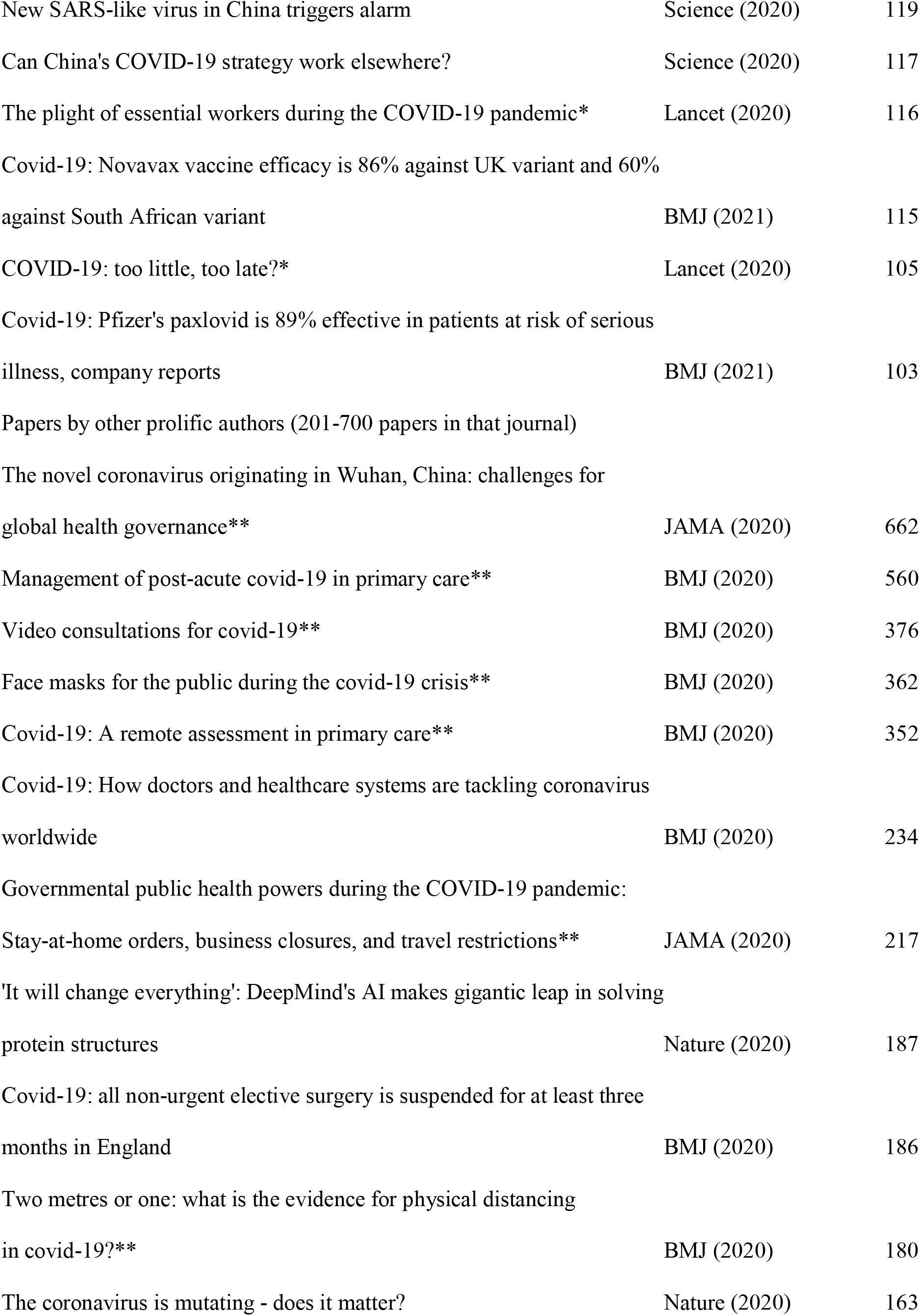
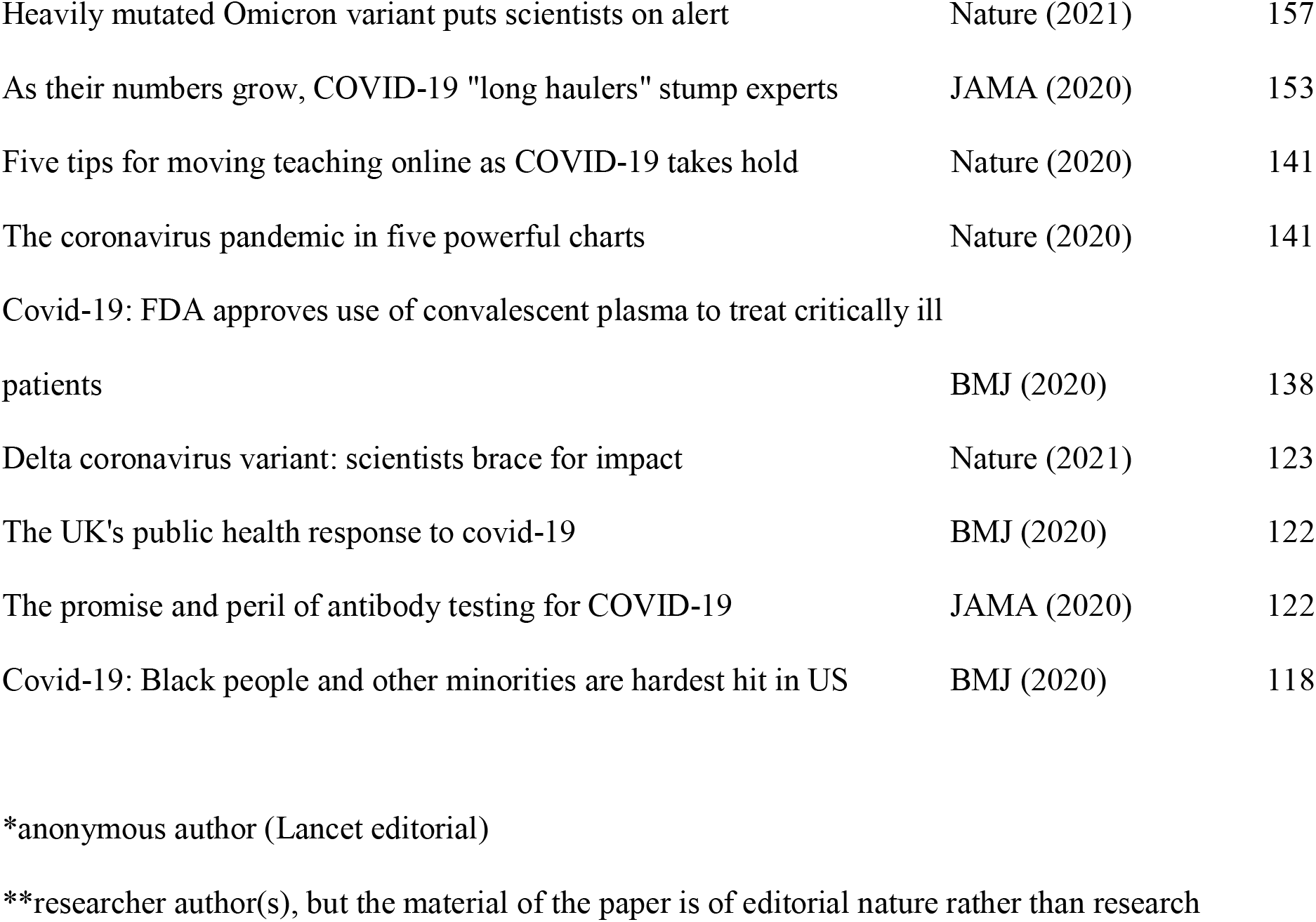
Highly-cited (>100 citations) journalistic papers published in 2020-2022 by non-researcher authors who have been prolific in their affiliated journals (>200 papers published in that journal)

34 of these 41 highly-cited papers were journalistic contributions by non-researcher authors. The other 7 were contributed by 2 researcher authors (with co-authors), however even these specific 7 papers were of editorial nature rather than research contributions. As shown in Table 3, most of the covered topics in the 41 highly-cited papers had public policy implications, including some of the most critical and momentous decisions for public health response, e.g. lockdowns and other measures taken by different governments and public health authorities. Many covered topics also pertained to situations where evidence was just emerging, e.g. the journalistic article was published promptly upon the press release of non-peer reviewed results from some studies or other preliminary observations.

### In-depth evaluation of the most massively prolific authors

29 individual authors were massively prolific, i.e. had published more than 700 papers in their main affiliated journal. 4 NEJM editors of Case Record items were excluded from further consideration, as it was unclear whether they should be considered authors of these pieces (as discussed above). For the remaining 25 individual authors, a search of potential additional Scopus ID author files showed that their main author ID files included 95% of the papers they had published in their affiliated journals with 2 exceptions. The median number of publications in their affiliated journal was 968 (range, 707 to 2302) and the median number of their total Scopus-indexed publications during their career to-date was 969 (range, 707 to 2302). The median number of publications in Scopus-indexed journals other than the one where they were massively prolific was only 4 and almost all their additional publications were also journalistic items. While two authors would qualify as prolific (>200 publications) for two different high impact journals (782 papers in Nature and 255 in Science; 790 papers in BMJ and 225 papers in Lancet), most massively prolific authors published entirely or almost entirely in only one of the assessed high impact journals during their careers.

Figure 1 shows the number of citations that these 25 authors had received during their career and the number of citations that they had received in papers published in their main affiliated journal. As shown, many of them were highly-cited with the median being 2293 citations for their total work (range, 135 to 22557, IQR 1303 to 5464). All 25 authors had received all or almost all their citations through the papers they had published in their main affiliated journal. The median number of citations to papers they had published in their main affiliated journal was also 2293 (range, 92 to 14138, IQR 1297 to 5179). 11 massively prolific authors had received zero Scopus citations to any work outside their main affiliated journal and for another 9 the only citations they had received outside their main affiliated journal were also to journalistic writings; another 4 authors had only 7-71 Scopus citations to their most-cited non-journalistic work and only 1 (an editor-in-chief) had highly-cited non-journalistic Scopus-indexed papers outside of their affiliated journal.

**Figure 1.**
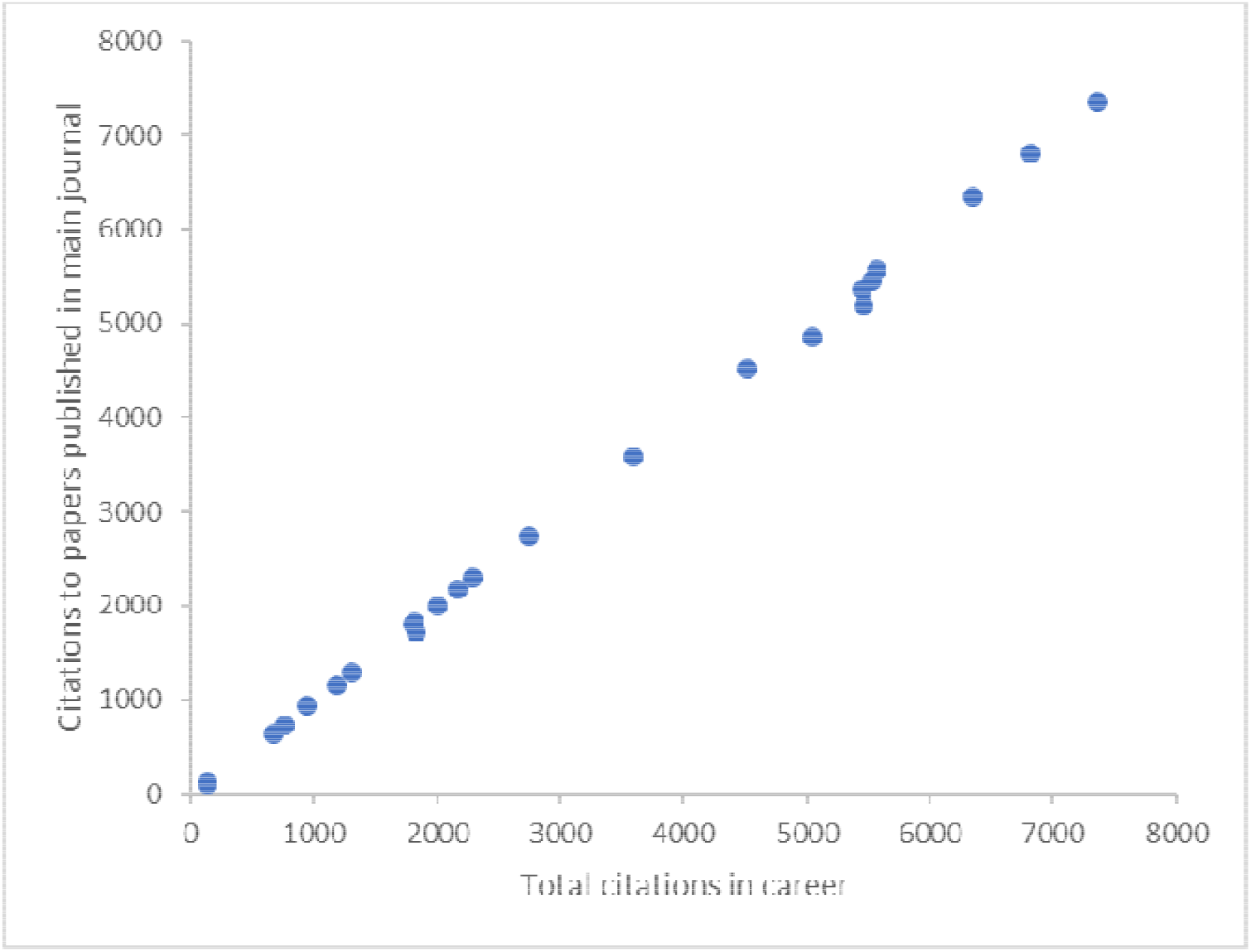
Total number of citations received during their career (horizontal axis) and total number of citations received to papers published in their main affiliated journal for the massively prolific authors with >700 publications in their main affiliated journal. Of the 25 eligible authors, one outlier is not shown.

The 25 authors had covered a wide range of hot, important topics in their most-cited journalistic articles including energy/climate change, addiction/behavior, gene therapy, global arsenic toxicity, junk DNA, diverse infectious diseases, 3-dimensional cultures, translational research, and opioid prescription abuse. All their recent (2020-2022) highly-cited papers (>100 citations) were on hot, rapidly emerging areas of COVID-19 (Table 3). The highest annual productivity was by an author who published 232 journalistic papers in BMJ in 2020.

Based on Google searches, 3 of the 25 massively prolific authors had obtained a PhD degree (in oceanography, pharmacology, and organic chemistry); however, the topics that they covered journalistically were typically not related to their PhD degree. At least two had an MD degree and another 7 has a Master’s degree in journalism or a related field. The other 13 seemed to have neither any doctoral degree (in any subject) nor a Master’s degree in journalism or in a related field. However, one cannot exclude that some education credentials were not disclosed/retrievable.

Among journal websites, BMJ provided systematic information on potential conflicts of interest for journalistic contributors. However, even in BMJ, of the 10 massively prolific authors, 3 had no entry for conflicts of interest, 4 replied in the online form that they had no conflicts of interest, 1 provided a vague statement that did not name the specific paying organizations and entities, 1 listed some specific organizations/entities but with no date, and 1 listed some specific organizations/entities and stated that the last update was in March 2019.

## DISCUSSION

The current analysis found many prolific authors who have published >200 papers each in at least one of the most prestigious journals of science or medicine at large. Almost all of them have published this work in a non-researcher capacity for a total of close to 70,000 journalistic publications in these highly sought publication venues. Nature, Science, and BMJ have the strongest representation of such prolific non-research authors. Many of these writings are very lengthy contributions (occasionally even longer than main original research articles) and Scopus characterizes over a third of the journalistic publications as full “articles” and another 11% as “short surveys”, while “editorial” is an uncommon characterization. Many of the published papers exert large influence in the scientific literature, as testified by large numbers of accrued citations. Almost all the highly-cited papers in 2020-2022 by such prolific authors were on COVID-19. In-depth evaluation of the 25 massively prolific authors, each with >700 publications in one of the most prestigious journals, shows that many of them are highly-cited; they have published little or nothing in the Scopus-indexed literature other than in their main affiliated journal; and they have written on diverse hot topics over the years. Very few of them have doctoral degrees in any subject matter, and a minority have a Master’s degree in journalism. Readily retrievable information on potential conflicts of interest of these authors in the journals where they massively publish is very scant.

The analyzed authors largely belong to the broad group of science journalism. Science journalism is an extremely important activity and it can have major benefits. Popularizing science can make it accessible to more non-specialist readers and to the general population. Science journalism also tries to effectively defend science in difficult times where anti-science movements abound. The cohort of analyzed authors includes several legendary figures with major, acknowledged and awarded contributions and with tremendous talent. Nevertheless, the breadth, popular outlook and fluidity of science writing is both a benefit and a risk. While a competent science writer may cover many fields, technical understanding of each field is challenging for an outsider. Most prolific authors had limited graduate training in science and only a minority got a degree specifically in journalism. They mostly learn the job iteratively, by experience. This may not be necessarily detrimental and there is debate on what is the best way to educate and form science writers (Ryan & Dunwoody, 1975; Dunwoody, 2004; Druschke et al., 2022; Hinnant & Lee-Rios, 2009; Jensen 2010). A legendary prolific science writer, David Jones (famous for his Daedalus column in Nature), jokingly described himself as “a fraud, charlatan and court jester in the palace of science” (source: https://www.chroniclelive.co.uk/news/north-east-news/death-former-newcastle-university-lecturer-13429028). More evidence is needed on best education and continuing education practices for science writers.

While talent may help in science writing, one wonders about the ability of even the most talented science writers to understand complex scientific fields. This becomes an even greater challenge for fields that are new and speculative with no or limited evidence, and for topics that are emerging, where even experts are mostly ignorant. The most-cited papers of the analyzed prolific authors targeted hot, and often controversial, topics. In 2020-2022 almost all of their highly-cited pertained to COVID-19 issues, many of which attracted heated debate, controversy, and reversals of evidence in the shifting sands of the pandemic (Tikkinen et al., 2020). Much research evidence is so methodologically problematic that it represents waste (Glasziou, Sanders & Hoffmann, 2020). It is unclear whether science writers can detect this efficiently in fields with massive production of papers (Ioannidis et al., 2021; Ioannidis et al., 2022). Importantly, journalistic papers in high impact scientific journals can be published within hours of having some new observation, preliminary data, or press release. Hence, these journalistic articles far outpace the publication timing of peer-reviewed scientific work. Therefore, these science writers may help shape strong opinions and guide positions or even policy pre-emptively. A perusal of the journal or personal sites of several of these science writers shows that many of them explicitly wish to focus on science policy, or even politics. Their views can be extremely influential. If well-informed or well-speculated, this is a great contribution, but if ill-informed or mis-conceived, this can create problems.

The influence of prolific science writers may vary enormously. This is shown by the variability in the citations to their work, and it should be acknowledged that only a minority of the papers they write attract large citations (Plomp, 1990). Regardless, variability in influence extends well beyond citations. The status and writing phenotypes of prolific authors may shape also how much their voice is heard. For example, an influential editor-in-chief may have more influence than an average news correspondent. Also some correspondents may have far more visibility, readability, immediacy, taste for controversial and hot topics, social media audience, and other reverberating features compared to others. Perceived importance, controversy, elite status (afforded by the publishing journal), and scale are factors that increase social media and media visibility and thus also wider impact (Htoo, Jin-Cheon, & Thelwall, 2022). Each major journal may also vary in the legacy of its editorial columns. For example, it has been found that Nature devotes more attention to internal science policy issues and Science more to the political influence of scientists (Waaijer, van Bochove, & van Eck, 2011).

Regardless, given the potential for major influence on both science, policy, and the community at large, science writers should be transparent about conflict of interest disclosures. This applies even more so to those who are the most prolific. The current analysis found a dearth of reported disclosures in the relevant journal sites. It was not analyzed whether conflicts of interest might have been disclosed in each single writing of these authors. Nevertheless, since these authors publish many hundreds of papers, it is essential to make full, updated lists of specific disclosures readily available. The few available disclosures suggest that some prolific authors may be paid by a large variety of sources, many of which may have direct or indirect ideological, political, business (pharma, big tech), or other agendas with financial repercussions. Furthermore, anonymity (e.g. papers signed by an editorial team rather than named individuals) should be discouraged. It is important to know who are the individuals penning influential commentary and what are their potential conflicts.

Some limitations of this analysis should be discussed. First, several of these prolific authors may also publish in non-scientific venues not indexed by Scopus, e.g. newspapers, social media, and political or science-oriented, technology, or general interest magazines. Therefore, the breadth of their influence may be much larger than what can be gleamed from their scientific journal corpus alone. However, publication in scientific venues carries a different level of seeming scientific authority. Second, the analysis focused on the extremely productive writers, but a much larger number of journalistic authors exist with fewer than 200 publications. Therefore, this analysis offers an obvious underestimate of the cumulative impact of the science writer community on the scientific journal literature. Third, the boundaries between the journalistic, editorial and research space may not be always sharply delineated and some individual authors may thrive in different spaces. However, for researchers it is notoriously difficult to have such a pervasive presence in a single journal; their work is typically spread across many publication venues. Fourth, other scientists have previously voiced concern about editors-in-chief publishing in their own journals (Scanff et al., 2021; Helgesson et al, 2022). Here, one should separate between original research and editorializing. It may be best to avoid publishing original research in one’s own journal during one’s editorship (Scanff et al., 2021), although this is not an absolute barrier. For editorializing activities, conversely, there is no felt restriction. If fact, editorials and other journalistic articles inflate impact factors: they do not count in the denominator of published articles, while their citations count in the numerator (Ioannidis & Thombs, 2019; Jain et al., 2021). Therefore, a perverse incentive may exist for editors to publish more journalistic pieces. The volume, features, and citation impact of editorial material varied a lot across journals (van Leeuwen et al., 2013).

A journal with a large, non-technical magazine section may engage more heavily in major debates, shaping science, action, and advocacy. However, are non-researcher writers the best choice to cover wide-ranging topics? Professional editors are typically not technical experts (Swidler, 2012; Editorial Nature Chemical Biology, 2011) and the same applies to recruited freelance science writers. One alternative option is for professional editors and science writers to co-author their pieces with knowledgeable subject-matter specialists. Alternatively, professional editors may offer more space to knowledgeable field-specific specialists and diminish their own presence and the presence of science writers. A third option is to include some supporting original data and systematically collected and appraised evidence in journalistic articles. This would require better training of science writers in rigorous meta-research methods (Ioannidis et al., 2015) and/or pairing with meta-researchers in co-authorship.

Eventually, prolific science writing in major scientific journals can be very influential. It is important to create a future research agenda to understand how to optimize this science writing corpus and how to ensure that these tens of thousands of produced papers age well both for science and for the general public. Moreover, given that survey data suggest that for scientific communication practitioners, 4 elements of trust are essential – competence, integrity, benevolence and openness (Besley, Lee & Pressgrove, 2021) – routine availability of disclosures for influential science writers would strengthen trust.

